# Tissue Resolved, Gene Structure Refined Equine Transcriptome

**DOI:** 10.1101/061952

**Authors:** T. A. Mansour, E. Y. Scott, C. J. Finno, R. R. Bellone, M. J. Mienaltowski, M. C. Penedo, P. J. Ross, S. J. Valberg, J. D. Murray, C. T. Brown

**Author notes:** Both authors contributed equally to this manuscript (T. A. Mansour, E. Y. Scott).

## Abstract

**Background:** Transcriptome interpretation relies on a good-quality reference transcriptome for accurate quantification of gene expression as well as functional analysis of genetic variants. The current annotation of the horse genome lacks the specificity and sensitivity necessary to assess gene expression especially at the isoform level, and suffers from insufficient annotation of untranslated regions (UTR). We built an annotation pipeline for horse and used it to integrate 1.9 billion reads from multiple RNA-seq data sets into a new refined transcriptome.

**Results:** This equine transcriptome integrates eight different tissues from 59 individuals and improves gene structure and isoform resolution while providing considerable tissue-specific information. We utilized four levels of transcript filtration in our pipeline, aimed at producing several transcriptome versions that are suitable for different downstream analyses. Our most refined transcriptome includes 36,876 genes and 76,125 isoforms, with 6474 candidate transcriptional loci novel to the equine transcriptome.

**Conclusions:** We have employed a variety of descriptive statistics and figures that demonstrate the quality and content of the transcriptome. The equine transcriptomes that are provided by this pipeline show the best tissue-specific resolution of any equine transcriptome to date and can serve several types of downstream analyses.

## Introduction

Transcriptomics is rapidly evolving from a focus on novel gene identification to resolving structural gene details. The transcriptomes of better-studied organisms, such as Drosophila, mouse and human have been updated to accommodate for this transition [1–3]. However, for less well characterized animals, such as the horse, there is often only annotation of a single variant of a gene with insufficient annotation of multiple splice variants, UTR extensions and non-protein coding RNA. This lack of information can challenge subsequent differential gene expression analyses and functional studies. There have been several attempts to improve the equine transcriptome with single tissue transcriptomes from lamellar tissue [4] or peripheral blood mononuclear cells [5] and from pooled composites of various tissues [6, 7], however a broader effort defining and integrating many tissue-specific transcriptomes and obtaining the library depth and strand information required to capture gene complexity is still needed.

ENSEMBL and NCBI provide publically available annotations for several vertebrate genomes including horses [8]. Both underlying annotation pipelines integrate homology search and *ab initio* prediction however accurate UTR prediction and isoform recognition require species-specific transcriptional evidence [9, 10]. For this equine transcriptome, the transcriptional evidence provided by total RNA sequencing (RNA-seq) was the basis of our gene annotation. This approach permits more reliable discovery of novel genes and isoforms, extension of UTRs and the flexibility necessary to establish a balance between sensitivity and specificity of gene detection for downstream applications.

Our annotation integrates the benefits of increased depth in reads and strand-specificity, for some tissues, as well as using a range of tissues from many horses, which allows tissue-specific transcriptomes to be extracted. We have incorporated RNA-seq from a diverse set of 8 tissues ranging from the central nervous system (CNS), skin and skeletal muscle tissues in adults to the inner cell mass (ICM) and trophectoderm (TE) in embryonic tissues (Table 1). The diversity in age, sex and tissue of the samples included in our assembly supply the equine transcriptome with its best spatiotemporal resolution and most complete gene UTR definition to date.

**Table 1.** Sample and library preparations used as input for our equine transcriptome.

| Tissue | Library Preparation | Library Characteristics | #Samples | #Frag(M) | #bp(Gb) | Reference |
| --- | --- | --- | --- | --- | --- | --- |
| BrainStem | RiboRNA-depleted | PE100bp, stranded | 8* | 166.73 | 33.68 | Finno et al, 2016 |
| Cerebellum | RiboRNA-depleted | PE100bp, stranded | 12 | 411.48 | 82.3 | Scott et al, 2016 |
| Muscle | Poly-A capture | PE125bp, stranded | 12 | 301.94 | 76.08 |  |
| Retina | Poly-A captured | PE80bp unstranded | 2 | 20.3 | 3.28 | Bellone et al, 2013 |
| SpinalCord | RiboRNA-depleted | PE100bp, stranded | 16* | 403 | 81.4 | Finno et al, 2016 |
| Skin | Poly-A captured | PE80bp, unstranded | 2 | 18.54 | 3 | Holl et al, 2016 |
|  | Poly-A captured | SE80bp, unstranded | 2 | 16.57 | 1.34 | Holl et al, 2016 |
|  | Poly-A captured | SE95bp unstranded | 3 | 105.51 | 10.02 | Bellone et al, 2013 |
| Embryo ICM | Ovation RNA-seq | PE100bp, unstranded | 3 | 126.32 | 25.26 | Iqbal et al, 2014 |
|  | Ovation RNA-seq | SE100bp, unstranded | 3 | 115.21 | 11.52 | Iqbal et al, 2014 |
| Embryo TE | Ovation RNA-seq | PE100bp, unstranded | 3 | 129.84 | 25.96 | Iqbal et al, 2014 |
|  | Ovation RNA-seq | SE100bp, unstranded | 3 | 102.26 | 10.23 | Iqbal et al, 2014 |
| Total |  |  | 69 | 1917.7 | 364.07 |  |

\*Seven individuals had both brainstem and spinal cord tissue collected from them. Seven of the skin samples were taken from 5 individuals and one individual had both retina and skin sampled, bringing our total number of individuals to 59.

We recognize that availability of annotation criteria and integration of transcriptome data is paramount for systematically improving the equine transcriptome. Our goal is to encourage equine researchers to incorporate their transcriptomic data using our pipeline as the common annotation pipeline and our initial transcriptomes as a reference framework. We intend to continue improving equine gene annotation through better UTR definition, isoform splicing characterization and novel gene identification. The annotation presented in this paper will improve the gene structure definition in current databases and the accuracy of downstream analyses, including both differential gene expression analysis and genetic variant annotation in the horse.

## Results

### Overall Mapping Statistics and Gene Counts After Filtration

RNA-seq of 59 samples in 12 libraries from 8 different horse tissues provided 1.9 billion fragments and 364 Gb of sequence bases. A summary of the library preparation, number of horses per library and total number of fragments and bases provided by each tissue library can be found in Table 1. The overall average mapping rate for Tophat2 was ∼83% with concordance rates ranging from 29% to 89% (average 75%) for paired end libraries. Concordance rates seem to be affected by the type of library preparation, where polyA selected and strand-specific libraries have the best rates. Library specific mapping rates can be found in Supplementary Table 1. The initial Cufflinks assembly identified 117,019 genes/211,562 transcripts. After this initial analysis we applied four steps of filtration (Figure 1). Primary filtration of transcripts removed the likely pre-mRNA fragments by eliminating single exon transcripts that were present within introns or overlapping with exons of other multi-exon transcripts. After primary filtration there were 75,102 genes/162,261 transcripts. The second filter was implemented to remove isoforms likely to be experimental artifacts by excluding low abundant transcripts with less than 5% of total expression for their locus. The remaining 114,830 transcripts represented 75,375 genes. In the third filter, non-coding transcripts that lack any supporting evidence from NCBI or ENSEMBL annotations, non-horse gene models (“Other RefSeq” and “TransMap RefGene” UCSC tracks) or *ab initio* predictions (“Augustus”, “Geneid”, “Genscan” and “N-SCAN” UCSC gene prediction tracks) were excluded. This third filtered version of the transcriptome has 76,323 transcripts in 37,062 genes. The last filter was for removing likely erroneous transcripts. The mtDNA in mammals is known for gene overlapping and polycistronic expression [11], permitting inaccurate prediction of mitochondrial transcripts by Cufflinks; we therefore excluded the mitochondrial contigs from our filtered assembly. Also, short transcripts less than 201bp (192 transcripts in 184 genes) were removed because they are more likely to represent repetitive sequences or incomplete gene fragments. Once erroneous transcripts were removed, our final refined version of the transcriptome contained 36,876 genes (76,125 transcripts) including 15,343 single exon transcripts, 8,808 two-exon transcripts, and 51,974 transcripts with three or more exons. A version of our refined transcriptome that is merged with the NCBI and ENSEMBL annotations, with redundant transcripts removed, is also available. This is the most comprehensive product of our pipeline and is valuable for differential gene expression analysis in tissues other than those provided in our assembly. Summary statistics including N50, number of genes and Mb and average length of fragment for all six versions of the transcriptome can be found in Supplementary Table 2.

**Figure. 1.**
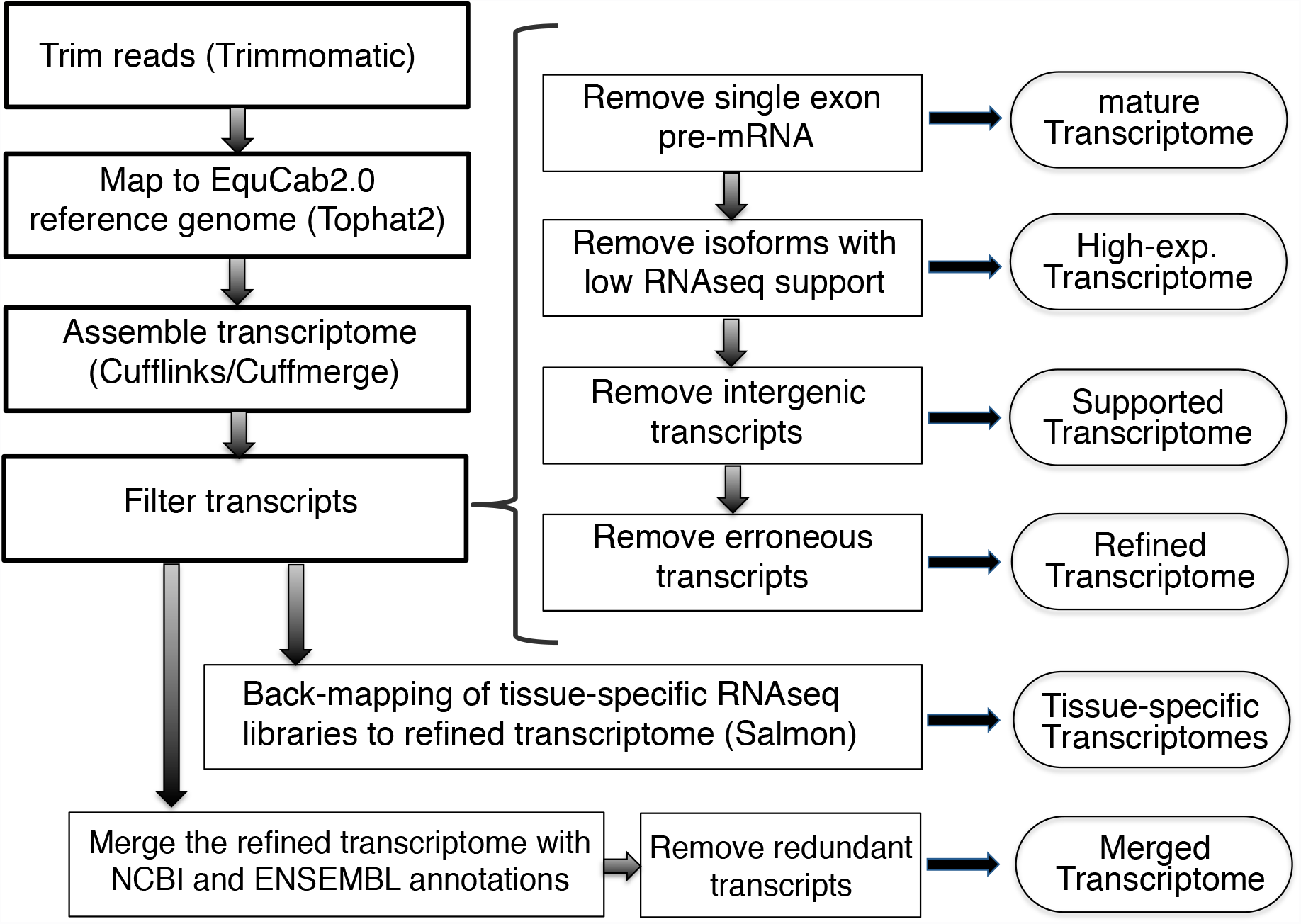
An outline of the workflow used to generate each version of the transcriptome. Transcriptome products are in ovals. Programs used to perform various steps are indicated in parentheses. All transcriptome versions and the pipeline scripts are publically available.

### Comparison between Our Transcriptome and Currently Available Equine Transcriptomes

We performed a comparison between our transcriptome and gene models from NCBI,ENSEMBL and two published equine transcriptomes, that we refer to as Hestand [7] and ISME [5] (Table 2 and Supplementary Table 3). In our comparisons, transcripts sharing one or more splice junctions are considered similar but only those with identical intron chains are matching. The comparison shows that the matching transcripts between our refined transcriptome and NCBI annotation are greater than 2.5-fold in number those matching the ENSEMBL annotation. However the highest number of matching transcripts occurred with the ISME transcriptome with 12,849 transcripts (Figure 2A). About 50% of the refined transcripts have a similar match in all the public transcriptomes.Evidence of improvements to the annotation of genes with a similar match to other assemblies can be found in genes such as *MUTYH,* where the three major isoforms annotated in humans [12] are now distinguishable in the horse (Figure 2B). The gene *CYP7A1* is another example where a novel first exon has been annotated and extended in our version of the transcriptome [13] (Figure 2C). About 20% and 28% of the refined transcripts are novel when compared to NCBI and ENSEMBL annotations respectively. Combined, there are 22,641 transcripts in candidate novel loci. Our approach of applying four successive steps of filtration strictly qualifies our novel isoforms as transcripts with ORFs or exonic overlap with candidate gene models. Mainly, novel transcripts contained within introns of other genes were excluded to avoid artifacts from retained intronic reads, common in rRNA depleted libraries. Using NCBI as a reference for comparison, our novel transcripts from the refined transcriptome have no bias towards any particular chromosome after accounting for chromosome size (Supplementary Figure 1). In order to calculate the gene and isoform detectability of our transcriptome compared to current annotation, we calculated sensitivity and specificity [14] between our transcriptome and a reference and found that, using NCBI as the reference, our transcriptome had a 78.8% sensitivity and 23.8% specificity at the base level and a 32% sensitivity and 21.1% specificity at the locus level. Detailed pairwise assessment for all equine annotations can be found in Supplementary Table 4. We developed a statistic to assess the conflict between different assemblies, termed “complex loci”, which refer to the loci that represent one gene locus in one transcriptome and two or more gene loci in another. Our transcriptome has 1355 and 997 transcripts that were considered complex loci between our transcriptome and NCBI and ENSEMBL, respectively. The Hestand transcriptome, however, has fewer with 660 and 798 complex loci when compared to NCBI and ENSEMBL, respectively. The ISME transcriptome has substantially more, with 1546 and 1226 complex loci when compared to NCBI and ENSEMBL, respectively.

**Table 2.** Comparison of current public equine annotations to six versions of our transcriptome (bolded and outline in red) in terms of gene numbers and composition

|  | **Unfiltered** | **Mature** | **High-exp** | **Supported** | **Refined** | **Merged** | Hestand | ISME | NCBI | ENSEMBL |
| --- | --- | --- | --- | --- | --- | --- | --- | --- | --- | --- |
| Genes (super-loci) | 117019 | 75102 | 75375 | 37062 | 36876 | 47760 | 56495 | 42654 | 24342 | 26962 |
| Transcripts | 211562 | 162261 | 114830 | 76323 | 76125 | 121997 | 68594 | 285538 | 43417 | 29196 |
| Multi-transcript loci | 17136 | 15430 | 14602 | 14511 | 14505 | 17835 | 8465 | 23833 | 7257 | 1592 |
| Multi-exon transcripts | 108985 | 108985 | 61570 | 60839 | 60782 | 97654 | 30949 | 259556 | 39272 | 19805 |
| Redundant transcripts | 0 | 0 | 0 | 0 | 0 | 2 | 3 | 12578 | 141 | 79 |
| Unstranded transcripts | 46928 | 35881 | 35872 | 6676 | 6618 | 5705 | 37673 | 4732 | 0 | 0 |
| Single exon transcripts | 102577 | 53276 | 53260 | 15484 | 15343 | 24341 | 37642 | 13404 | 3723 | 6862 |
| Two exons transcripts | 11114 | 11114 | 9449 | 8857 | 8808 | 11350 | 4410 | 13092 | 2425 | 2972 |
| Many exons transcripts | 97871 | 97871 | 52121 | 51982 | 51974 | 86306 | 26542 | 259042 | 37269 | 19362 |

**Table 3.** Tissue-specific splicing rate as calculated by Cuffcompare, with relevant number of multi-exonic transcripts and multi-transcript loci per tissue.

|  | Embryo<br>ICM | Embryo<br>TE | Skin | Brainstem | Cerebellum | Retina | Spinal<br>cord | Muscle |
| --- | --- | --- | --- | --- | --- | --- | --- | --- |
| Genes | 33998 | 32050 | 30003 | 34792 | 36139 | 26733 | 34980 | 29549 |
| transcripts | 57400 | 54424 | 51995 | 62993 | 66364 | 47095 | 66001 | 52000 |
| multi-exon<br>transcripts | 44069 | 42433 | 42432 | 49346 | 51640 | 39420 | 52175 | 42483 |
| multi-transcript<br>loci | 11938 | 11461 | 11797 | 13066 | 13334 | 10866 | 13352 | 11560 |
| Splicing<br>rate | 1.7 | 1.7 | 1.7 | 1.8 | 1.8 | 1.8 | 1.9 | 1.8 |

**Figure. 2.**
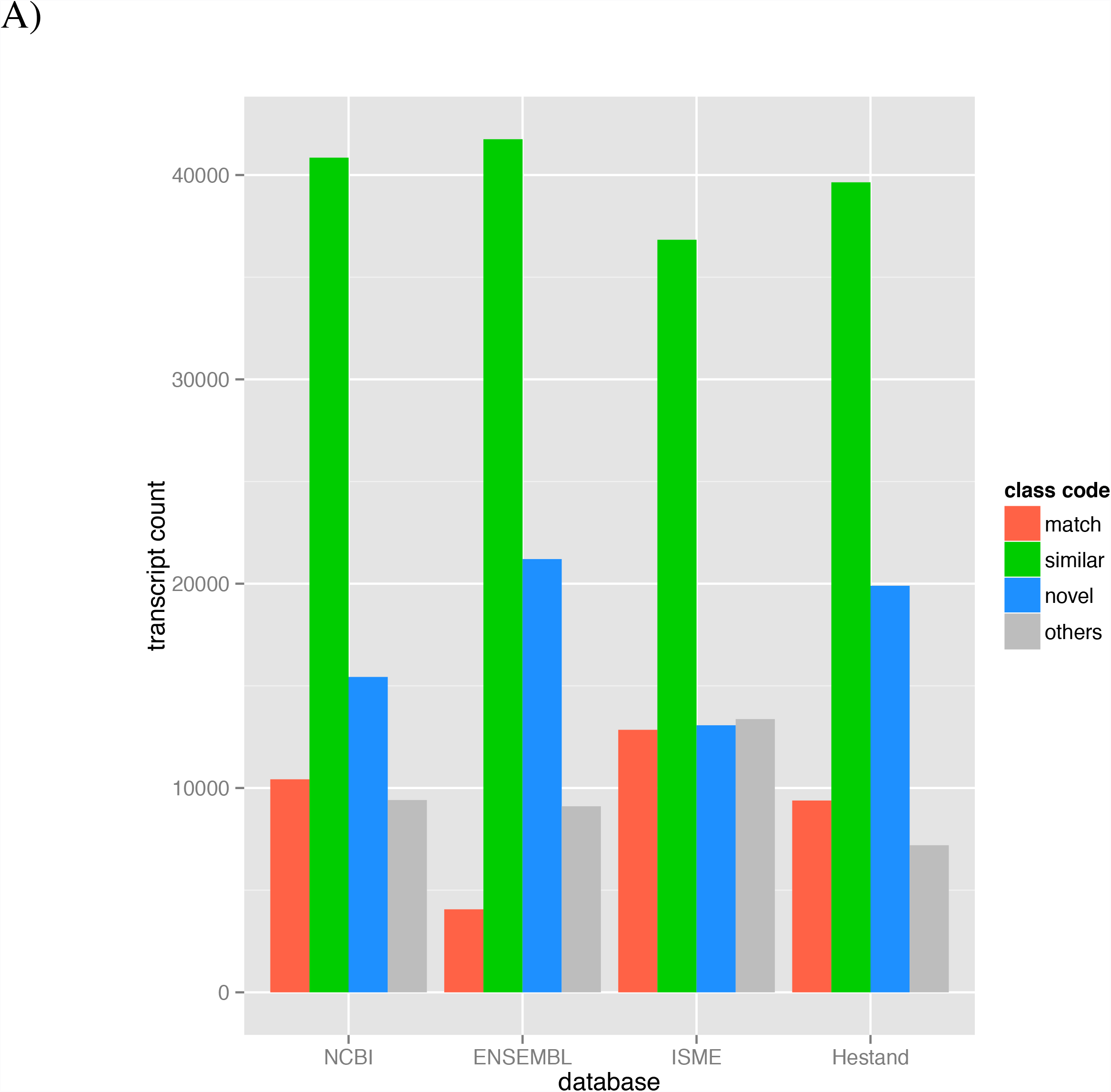
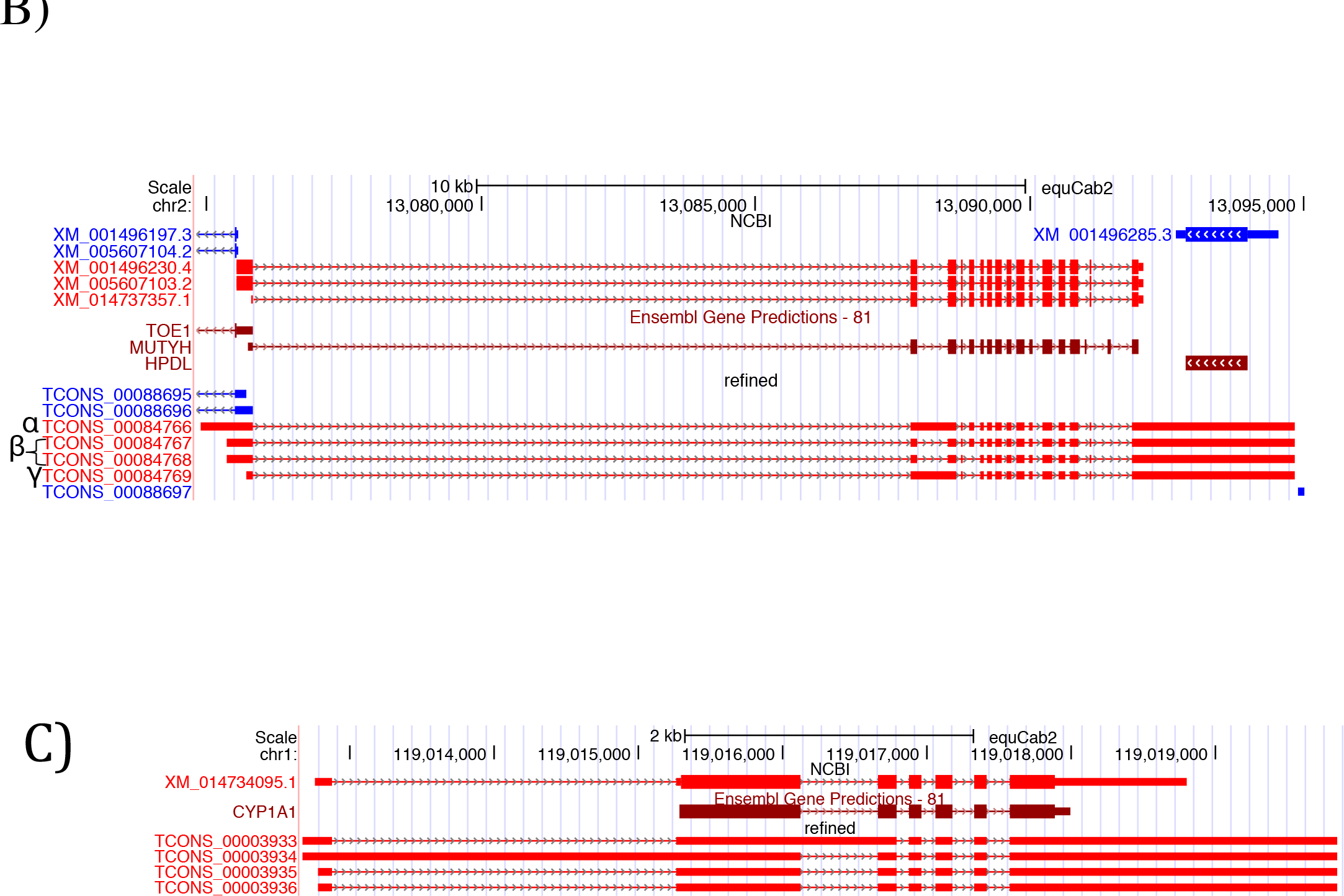
Comparison of our refined transcriptome to current equine annotations. (A) Our refined transcriptome compared to current annotations. (B) The annotation of *MUTYH* in the refined version of the transcriptome shows the addition of several isoforms, α, β, and γ, as seen in the human, of *MUTYH*. (C) The gene annotation of *CYP7A1* in the refined transcriptome also shows the inclusion of an extended alternative first exon not seen in other species.

### UTR extension

To test the effect of the new assembly on the UTRs of known genes, we identified the protein coding isoforms sharing the exact intron chain with NCBI isoforms, which yielded 9736 isoforms from 7419 genes. The difference in the total length of each transcript was then calculated and we found that we extended the length of 8899 isoforms (6817 genes) by 29.7 Mb in total. 831 isoforms (718 genes) lost 0.3 Mb in total with an average of 0.4 kb per isoform, while 6 isoforms did not change.

### Gene and Isoform Distinctions between Tissue-Specific Transcriptomes

We selected genes with high expression (a sum of TPMs across all tissues above 200) and substantial expression differences across tissues (a standard deviation above 200). Unsupervised hierarchical clustering grouped genes that may be co-expressed as well as illustrating the relationship between the tissue-specific transcriptomes. As expected, the transcriptomes from the three central nervous system (CNS) tissues clustered together, as did the two embryonic tissues, with the skin and skeletal muscle furthest from these clusters (Figure 3A). Blocks of genes showing uniquely high expression in a given tissue were further annotated with NCBI gene names and then summarized with Panther biological processes annotations. The top two Panther pathways (lowest p-values) for each of these gene blocks are reported in the text below, with the corresponding p-values (p) and fold-enrichment vales (FE). The full Panther annotation tables are detailed in Supplementary Table 5. The CNS cluster contained overrepresented processes regarding brain function and development: nervous system development (p=2.10E-06, FE=7.12) and neurogenesis (p=9.36E-05, FE=7.73). The retina contained processes consisting of photoreception and visual perception: phototransduction (p=3.52E-08, FE=80.75) and visual perception (p=3.69E-18, FE=37.64). The skeletal muscle encompassed genes pertaining to muscle physiology and assembly: muscle contraction (p=1.48E-27, FE =57.58) and myofibril assembly (p=6.05E-11, FE=72.8). The embryonic tissues have the most general processes assigned to their distinct clusters: translation (p=1.15E-11, FE=16.35) and peptide biosynthetic processes (p=1.95E-11, FE=15.8). And finally, the skin consisted of processes concerning epithelial organization and production: intermediate filament organization (p=1.69E-07, FE > 100) and skin development (p=1.89E-09, FE=22.49).

**Figure. 3.**
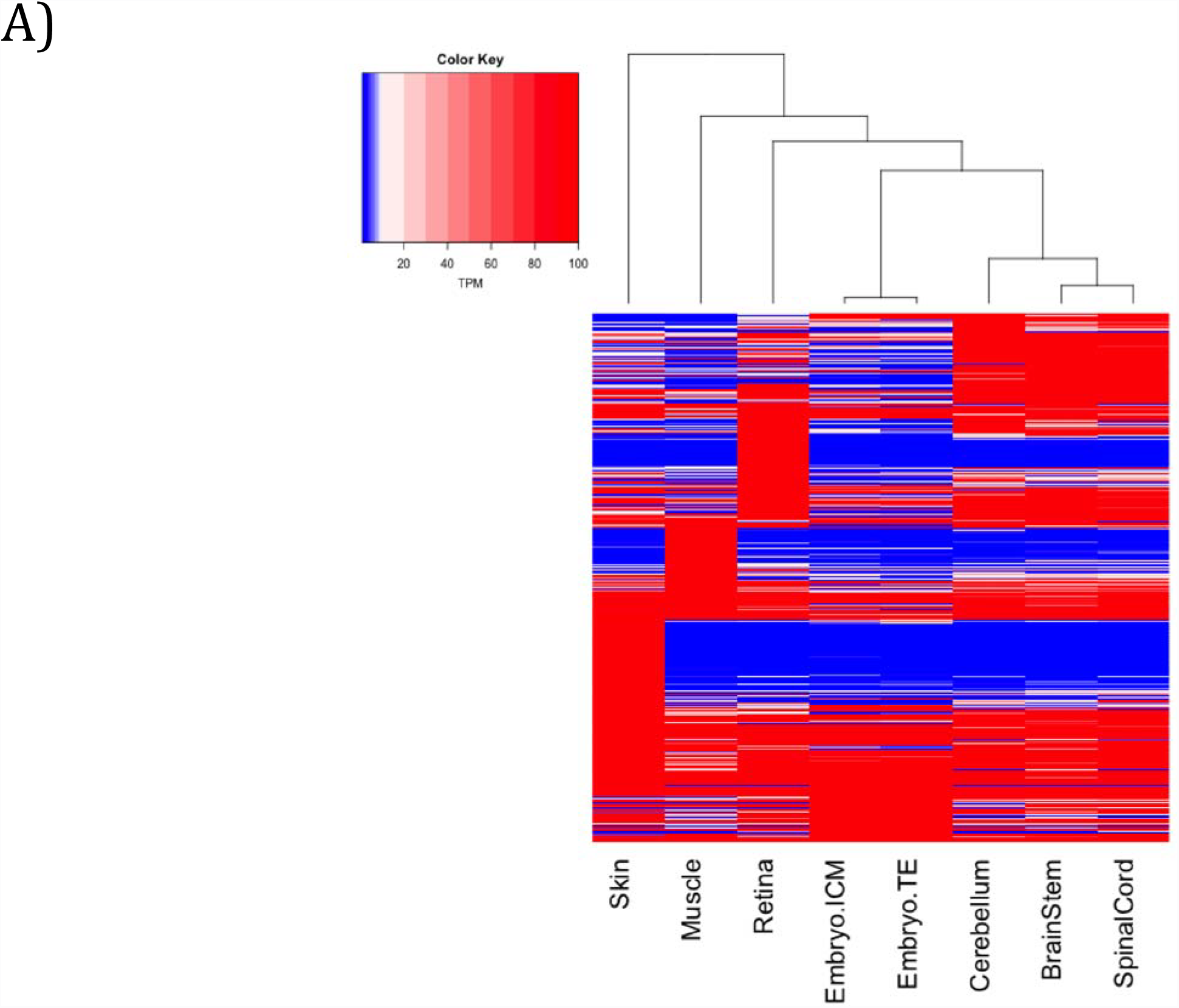
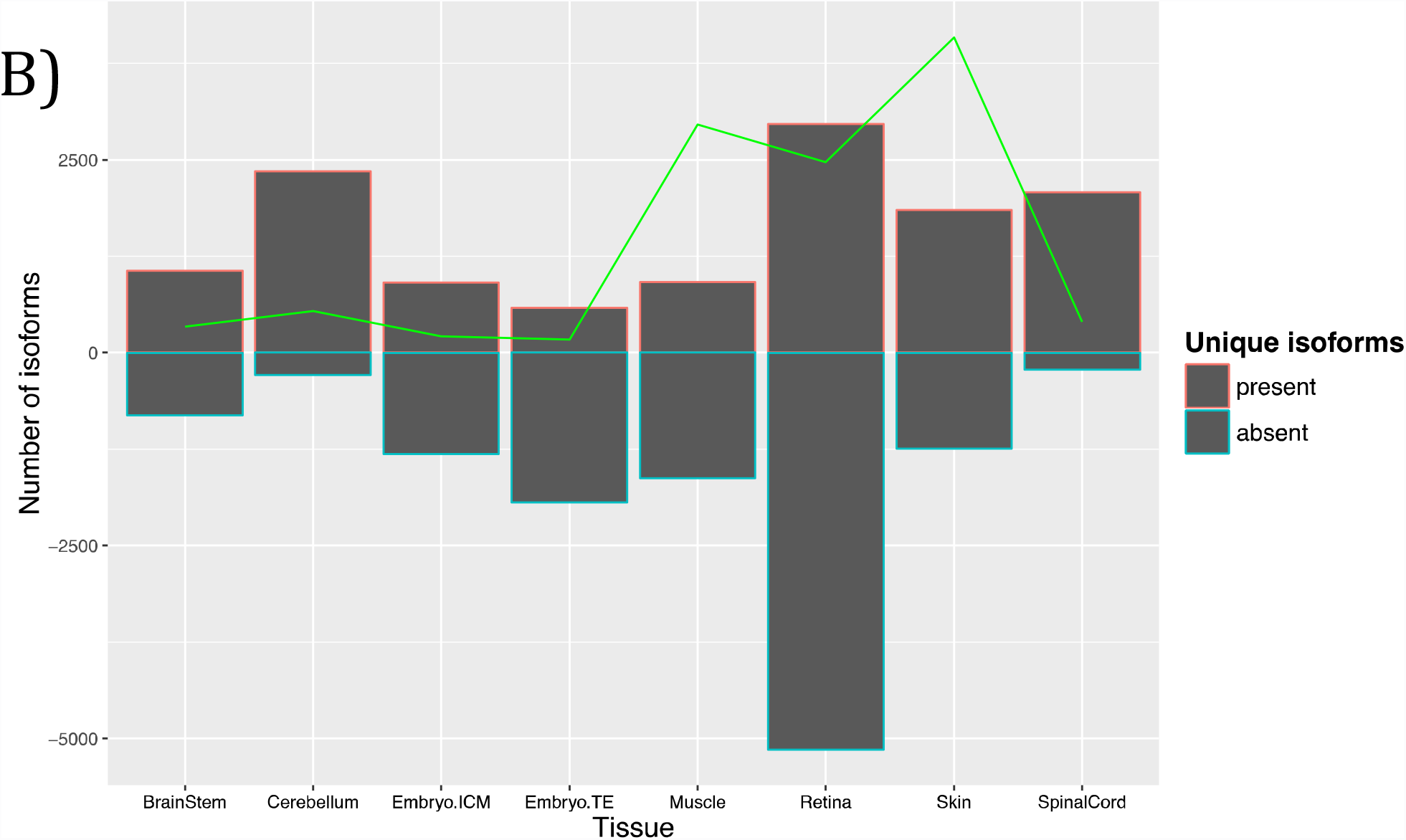
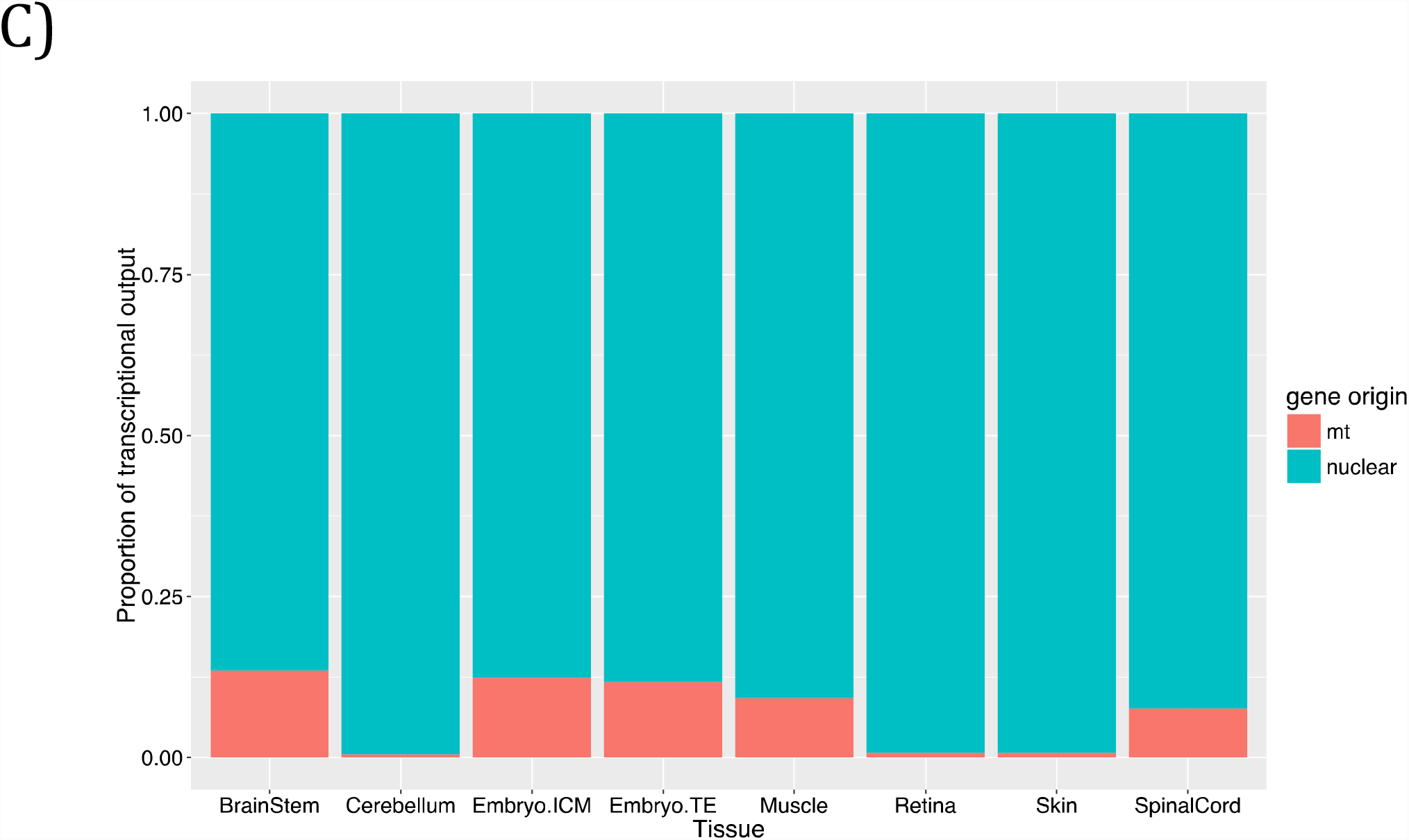
Tissue-specific gene and isoform composition of the transcriptome. (A) A heatmap of genes with high expression and substantial expression differences across tissues. (B) A bar graph showing isoforms uniquely present (the bar outlined in red above the x-axis) or solely absent (the blue outlined bars extending below the x-axis). The green trendline corresponds to the cumulative TPM of the uniquely present transcripts. (C) A stacked bar graph showing the transcription percentage of mitochondrial genes versus nuclear encoded genes.

When attention is given to the isoforms showing unique presence or sole absence in a tissue, the retina and cerebellum possess the most isoforms that are uniquely present, with the retina also containing the largest amount of solely absent isoforms (Figure 3B). The uniquely present transcripts in the retina have Panther annotations of visual perception (p=2.96E-23, FE=7.47) and photoreceptor cell differentiation (p=1.59E-09, FE=7.56) and in the cerebellum nervous system development (p=3.08E-09, FE=1.85) and generation of neurons (p=1.72E-08, FE=2.02). Full Panther annotation tables of transcripts uniquely present in tissues can be found in Supplementary Table 6. The transcripts solely absent from the retina pertain mainly to positive regulation of DNA replication (p=2.49E-03, FE=3.53) and anatomical structure development (p=2.72E-19, FE=1.48). Full Panther annotation tables of transcripts solely absent in tissues can be found in Supplementary Table 7. Utility of these isoforms, in terms of expression, is strongest in the skin, retina, skeletal muscle and to a small extent the cerebellum (Figure 3B). Despite these differences in unique isoforms, multi-exons transcripts and multi-transcript loci, the splicing rate across tissues, as calculated by Cuffcompare [15], ranges from 1.7 to 1.9 (Table 3).

Nuclear coding versus mitochondrial encoded genes were parsed out per tissue to determine how much of the sequencing resources are allocated to genes of the mitochondria (Figure 3C), with the conclusion that the brainstem, spinal cord, embryonic tissues and skeletal muscle exhibit the largest proportions of transcriptional output devoted to mitochondrial genes.

### Classification and Annotation of Novel Genes

In total there were 22,640 novel transcripts, with varying levels of support from current equine annotations (Figure 4A). Classification of these candidate novel genes was necessary to better describe our novel gene identification and aid in interpreting the degree of support available to each category. Three categories of novel genes based upon the supportive evidence within and across species were made with each successive category being less supported by equine or orthologous gene models. Our first category of novel genes contains those missing from NCBI and/or ENSEMBL annotations, but supported by either NCBI, ENSEMBLE, Hestand or ISME annotations (Category I). The second group of novel genes were novel to all public equine annotations, but conserved by means of orthologous gene similarity or supported by possible gene prediction (Category II). The third category of novel genes were unsupported by any candidate gene models, but had an ORF (Category III). Category I has a total of 8459 transcripts, with 2/3 of these transcripts novel to the ENSEMBL annotation (Supplementary Figure 2). Another 1849 transcripts in this category are absent in both NCBI and ENSEMBL annotations, yet supported by Hestand or ISME annotations. Homology with the SWISS-prot database identified at least one significant (p<1E-10) hit for almost half the transcripts in this Category I (Supplementary Table 8). The second category has 7494 transcripts that – unless on the opposite strand - do not overlap with known gene models in public annotations. Annotation of these transcripts was performed partially by testing overlap with non-horse gene models and also by homology search. Only 16% of these transcripts have significant hits against the SWISS-prot database (Supplementary Table 8). The third category of novel genes includes 6687 transcripts with an ORF as the only functional support for these transcripts. The first category of novel genes shows the most diverse distribution of exon numbers comprising the genes (ranging from 1 to 28), whereas the unsupported genes contained mainly single exon genes (Figure 4B). The expression analysis of the three novel gene categories shows a clear reduction of cumulative expression from category I to III. There was also an obvious tissue-specific pattern in the expression of novel genes. Supported novel genes (Category I) had the highest expression in the cerebellum. However, when looking at only the second category of novel genes, the embryo contributed the highest expression of novel transcripts. Category III novel transcripts mainly consisted of single exon transcripts and showed similarly low expression across all tissues (Figure 4C).

**Figure. 4.**
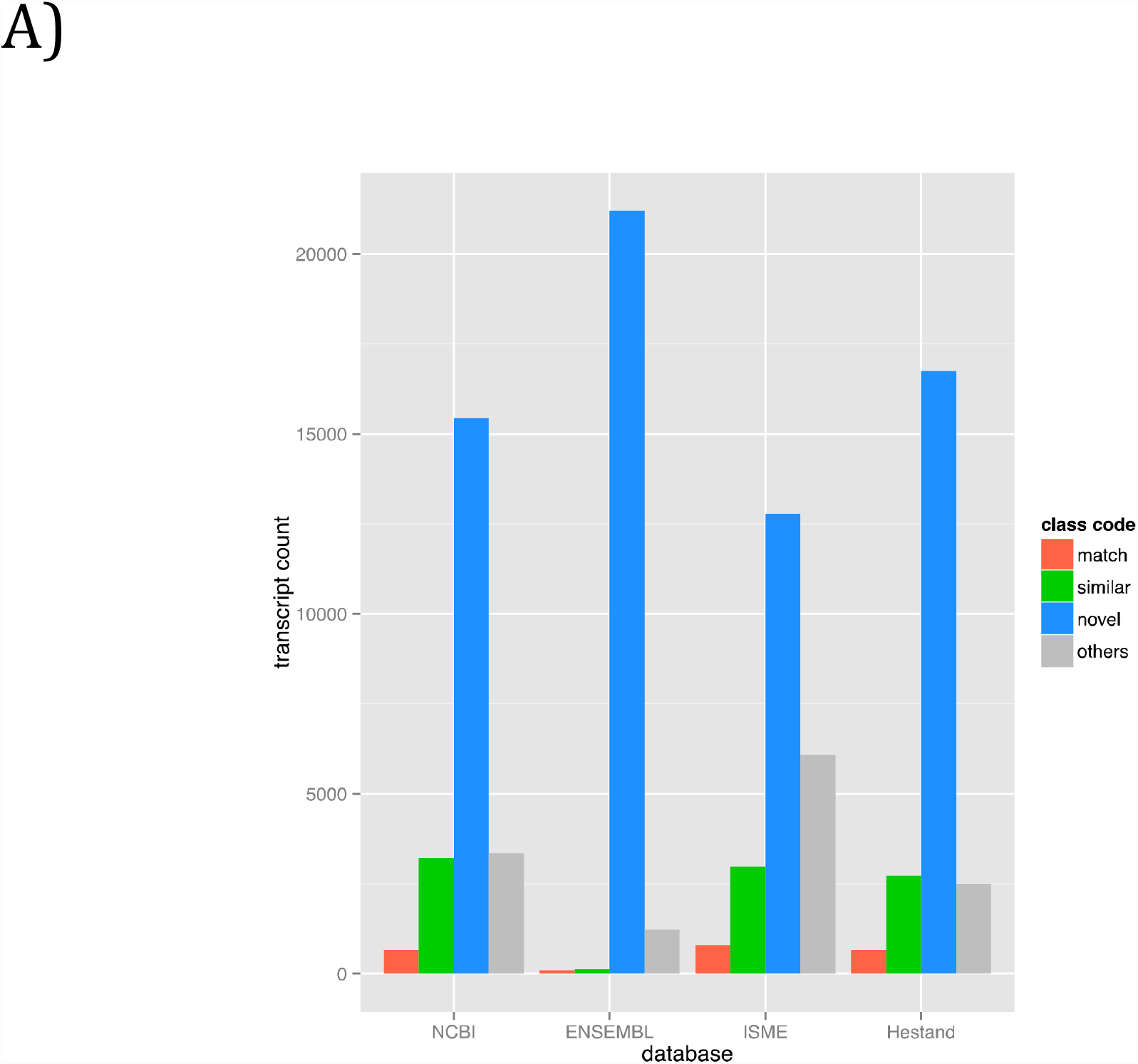
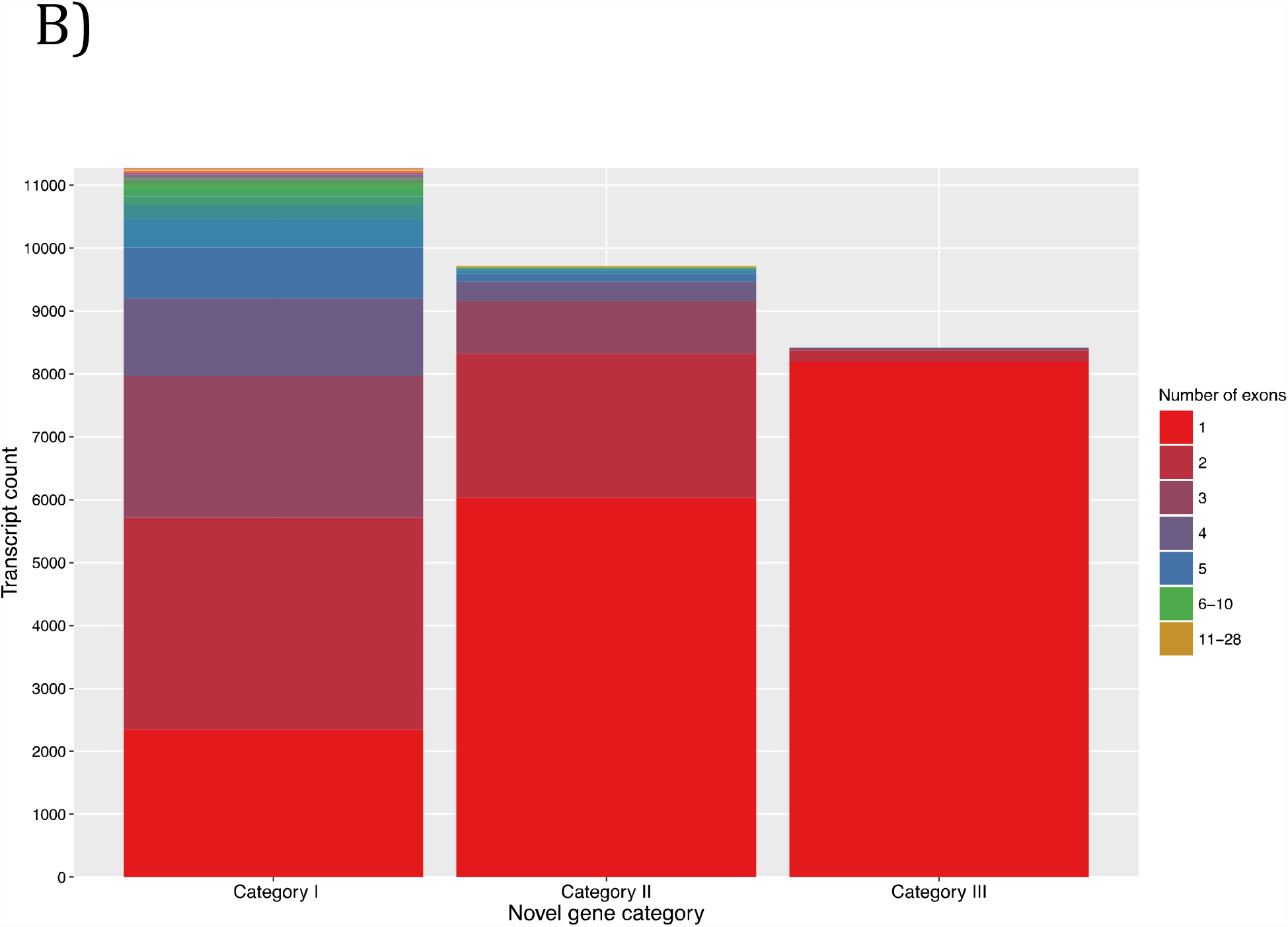
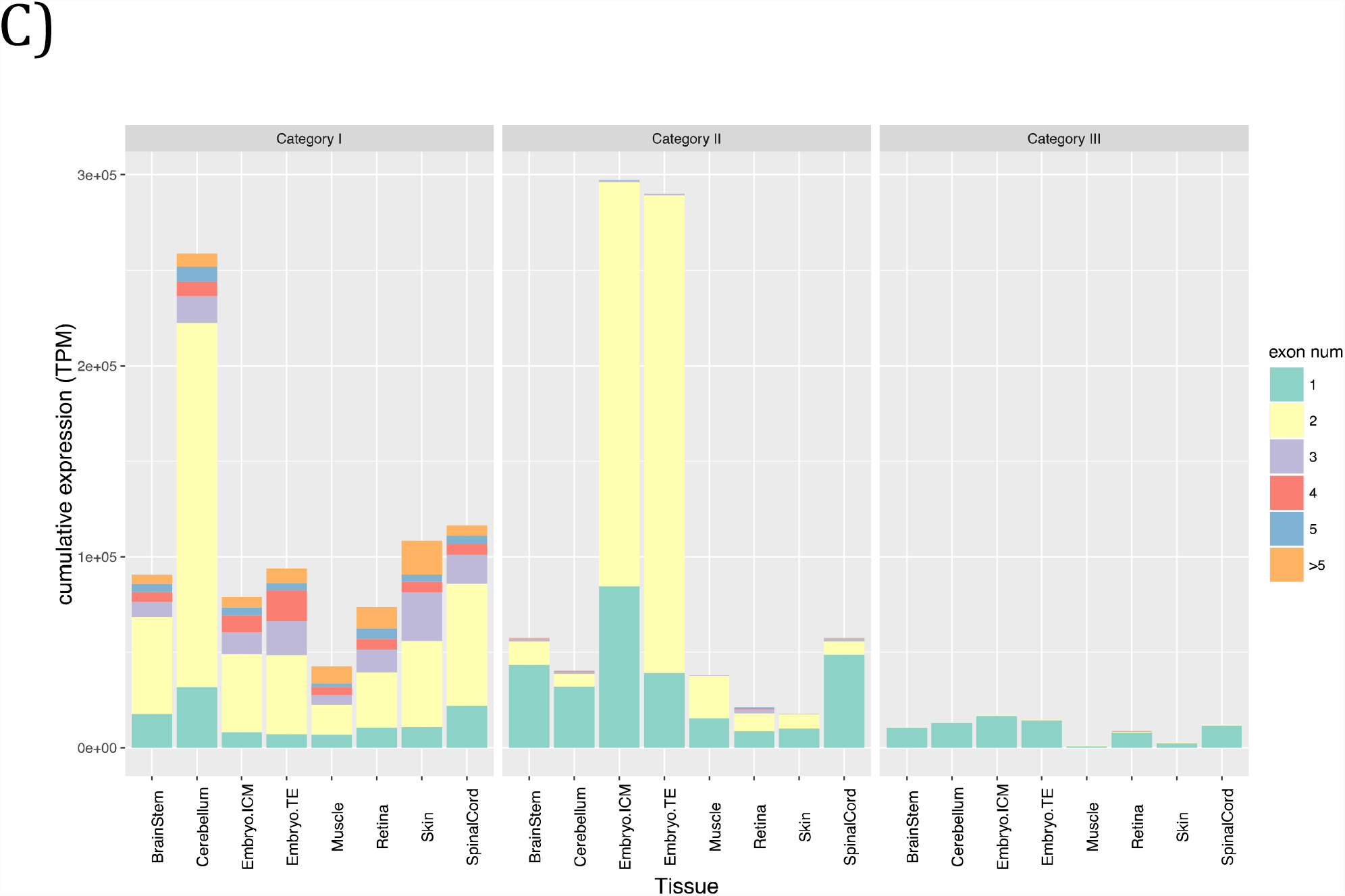
Novel gene analysis and classification. (A) A bar graph showing the comparison of all the novel genes against the current equine annotations. The three categories of novel genes were supported novel genes (Category I), unsupported, but conserved, novel genes (Category II) and the unsupported, un-conserved, but novel genes with an ORF (Category III). (B) A stacked bar graph of transcript counts with all three categories of novel genes showing exonic composition and (C) their cumulative TPM in a tissue specific manner.

## Discussion

Using RNA-seq from fifty-nine horses across eight tissues has allowed us to capture transcriptome complexity and provide spatial resolution in terms of tissue-specificity in manner that exceeds any current equine annotations. Our descriptive statistics and accessible pipeline make this project open to modifications and further integration of transcriptomes.

RNA-seq based transcriptomes are prone to false inflation of gene numbers for several reasons. Technical limitations such as limited sequencing read length and amplification errors, false splicing events, and assembler deficiencies are among several reasons of misassembly. Pervasive transcription is another predominant source of such inflation [16–18]. Some types of sequencing libraries increase the problem as well; for example rRNA depletion inflates the assembly with primary transcripts and false isoforms exhibiting intronic retention [19]. Our pipeline takes these factors into account and runs unguided by previous transcriptome annotations with several transcript filtration steps that reduce inclusion of inaccurate transcripts, while retaining the sensitivity for novel transcript detection. The effect of this procedure can be seen by comparing the gene numbers between our initial unfiltered and final filtered transcriptomes, where gene inflation was reduced by 68% (Table 2) and our final refined transcriptome contained 36,876 genes and 76,125 isoforms.

Although not indicative of transcriptome quality, we calculated specificity, as a measure of difference between our transcriptome and other annotations, and sensitivity, which indicates how our transcriptome covers another annotation. These parameters demonstrate that our aggressive filtering does sacrifice sensitivity at the locus level only by a margin of approximately 5%, and increases our specificity often by more than 10%, relative to NCBI and ENSEMBL (Supplementary Table 4). We have a comparable sensitivity to the Hestand transcriptome, which could be explained by adopting strict filtering approaches in both pipelines. However, the numbers of unstranded and multi-exon transcripts in the Hestand transcriptome relative to our refined version serve as the more discriminating statistics. We have approximately six fold less unstranded transcripts and more than double the multi-exon transcripts (Table 2). Regarding how our transcriptome compares to the other recent equine ISME assembly [5], which is ENSEMBL annotation guided, we have three times more matching transcripts to the ISME assembly than to the ENSEMBL annotation itself (Figure 2A), suggesting significant improvement made by the ISME annotation. However their improvements are impaired by false inflation in the number of genes identified due to presenting most of the transcripts in two copies representing the forward and reverse strands. This inflation of the ISME annotation can explain why it has different statistics from Hestand as well as our new transcriptome. Hestand et al (2015) also observed a bias towards single exon genes, which represented approximately 55% of their whole transcriptome [7]. Our native assembly identified similar percentage of single exon transcripts, however those numbers went down to 20% after filtration of single exon pre-mRNA. Our statistic, complex loci, also highlights a level of sensitivity as well as an area for further investigation in our transcriptome. We have more than two times more complex loci, using NCBI as a reference, than Hestand. The inflated ISME complex loci numbers could be attributed to the double reporting of their transcripts. Awareness of these complex loci allows for refinement of transcriptome-wide gene structure, while a pipeline to appropriately process these loci has yet to be established. The evaluation of these alternative descriptive statistics permits comparisons with transcriptomes that have pipeline-specific limitations.

Accurate identification of UTRs is often difficult for *ab initio* programs and requires sufficient support of transcription evidence. Our integrative analysis of several tissues using different library preps enabled us to achieve unpreceded extension of equine transcripts’ UTRs by an average of 3.3 kb per transcript. Indeed high coverage of CNS tissues in our analysis was an important factor as reported previously with several transcriptomes [20], [21] and [22]. Further improvements to this transcriptome would include providing tissue-specific UTR lengths and allowing for a more clear depiction of differences in gene structure between tissues. The improved UTR structure provided by our transcriptome has already shown its utility in the horse community by defining isoform and gene boundaries of *MUTYH* and *TOE1* [23] as well as providing an alternative start exon for *CYP7A1* [13]

The distinct RNA-seq libraries from eight tissues,allow us to extract tissue-specific features regarding gene expression, isoform usage and mitochondrial gene expression. Tissue specificity, in terms of gene expression, was demonstrated on two levels, first as biases in exon usage, especially one and two exon gene expression in the embryo (Supplementary Table 3). Second, gene clustering and Panther annotation revealed inherent functions associated with each tissue (Figure 3A, Supplementary Table 5). The retina displayed the most uniquely present and absent isoforms, in agreement with other studies [25], and had correlated isoform annotations unique to the retina: visual perception and photoreceptor cell differentiation. The skeletal muscle was a tissue with a relatively low amount of unique tissue-specific isoforms, however, it shows utility of these isoforms with relatively high cumulative expression values (Figure 3B) as well as comparable splicing rates (Table 3). Three tissues: retina, skin, and embryo, had shorter read lengths and were not prepared as stranded libraries and thus these data may be artificially understated in terms of transcriptome complexity. In addition to the nuclear gene expression, the amount of transcription occurring from the mitochondrial chromosome can show how much of the sequencing resources are being allocated to genes of mitochondrial origin.Across the eight tissues, one would expect the tissue with the largest numbers of mitochondria to have the largest proportion of mitochondrial allocated transcriptional output [2]. Our data demonstrates these trends in the brainstem, skeletal muscle and spinal cord, however the cerebellum and retina do show an unexpectedly low mitochondrial gene load (Figure 3C). Further research establishing the relationship between the amount of mitochondria processed in a sample for RNA-seq and the resulting mitochondrial expression loads would be beneficial to understanding how much of the transcriptional output is dominated by an individual mitochondrion.

We identified 7494 candidate novel transcripts. These novel transcripts are selected based on having no overlap with genes in current equine annotation and authenticated by their protein-coding ability and/or overlap with aligned non-horse genes or *ab initio* gene predictions. Our novel transcripts have a diversity of coding exons in Category I and a particular expression bias in the embryonic tissues of Category II, in which a majority of these novel transcripts contain two exons (Figure 4C). The Category I novel transcripts highlight the deficient equine ENSEMBL annotation, the need to pool the databases to get the most transcriptome coverage and the ability of our transcriptome to capture the potentially rare novel gene models (Figure 4A). Despite the ORF requirement for Category III novel transcripts, there is an obvious enrichment for single exon transcripts and a marked reduction of total transcription level (Figure 4C), which is indicative of non-coding RNA. Our novel gene analysis also produced a category of novel transcripts that were removed due to not having ORFs and were presumed to represent noisy transcription relating to primary transcripts, repetitive elements, sequencing errors and genome-based errors. The collection of Category III and these excluded novel transcripts may represent a repository of non-annotated non-coding RNA, which is an area that needs further annotation in the horse genome.

This transcriptome assembly pipeline not only produces flexible incorporation of additional transcriptomes, it also provides several products regarding levels of transcript filtering and appropriateness for downstream analysis. The different extractable transcriptome versions include transcriptomes after each individual filter, with the final refined transcriptome containing only genes with complete ORFs and genes aligning with other non-horse genes or *ab initio* gene predictions. A version of our transcriptome merged with NCBI and ENSEMBL annotations achieves breadth not covered by our tissues. These transcriptomes as well as the pipeline to make each of these transcriptomes are publically available on our GitHub repository. By making the workflow public and easy to execute and manipulate, we aim to expand the spectrum of tissues embodying this transcriptome and eliminate biases in annotated genes and thus downstream differential gene analysis. Increasing and providing the option for tissue-specific transcriptomes, allows for more targeted and refined usage of a certain tissue’s transcriptome for differential gene analysis, resulting in downstream analysis of more significant differentially expressed genes. As stated in our overall goals of this project, we have provided a framework for further improving the equine transcriptome and produced an equine transcriptome that expands on current equine annotations in the manner of UTR extension, isoform detection and novel gene identification.

## Methods

### RNA-seq library preparation

A total of twelve RNA-seq libraries in 8 tissues from 59 individuals (20 female, 27 males and 12 embryos) were used to prepare our transcriptome. The brainstem, spinal cord and cerebellum were strand-specific 100 bp paired-end (PE) libraries. The skeletal muscle tissues were strand-specific PE125 bp libraries. A subset of the embryo ICM (3 samples) and TE (3 samples) were unstranded PE100 bp libraries, the other subset (3 ICM and 3 TE) consisted of single end (SE) 100 bp reads. The retina RNA-seq libraries were unstranded SE80 bp libraries. And the skin libraries were all unstranded and consisted of PE80 bp, SE80bp and SE95 bp reads. The brainstem, spinal cord and cerebellum RNA libraries were all rRNA-depleted, the skin, retina and skeletal muscle libraries were poly-A captured. The embryonic libraries were neither poly-A selected nor rRNA depleted, they were prepared with the Ovation^^®^^ RNA-seq System V2 (NuGEN, San Carlos, CA, USA), which aims to amplify mRNA as well as non-polyadenylated transcripts. Table 1 summarizes the tissue-specific RNA-seq library parameters.

### Trimming and Mapping of Reads

The Illumina adaptors as well as the reads were trimmed with the sliding window quality trimmer Trimmomatic [26] with a window size of 4 and a softer quality threshold of 2 [27]. Mapping of the trimmed reads was done with Tophat2 [28] to EquCab2.0, 2007

(ftp://hgdownload.cse.ucsc.edu/goldenPath/equCab2). Cufflinks [15] was used to assemble transcripts from the aligned RNA-Seq reads. Two cerebellar samples failed assembly due to computation limitations (8 CPUs, 250 Gb RAM and 7 days) and required digital normalization [29] to 200X coverage before mapping with Tophat2.

### Filtering Transcripts

Four categories of filters were used to remove likely pre-mRNA and artifactual transfrags, as summarized in Figure 1, resulting in six versions of the transcriptome. Primary transcript filtration was done using Cuffcompare [15] between our assembly and a version of our assembly containing only multi-exon transcripts and removing transcripts overlapping with intronic regions and with class codes “i”, “e” and “o”. The input trimmed RNA-seq reads were then back-mapped to the pre-mRNA free transcriptome using the quasi-mapping based software package Salmon [30]. While back-mapping, a second filtration step was implemented: low abundance transcripts in every locus were excluded with the lower threshold of a TPM (normalized read count standing for transcripts per million) less than 5% of the total TPM per locus. For the third filter, Transdecoder [31] was used to predict the ORFs and Cuffcompare [15] to determine any exonic overlap with any candidate gene locus using the class codes “j”, “o”, “x” and “c”. In the Transdecoder analysis, the longest open reading frames were extracted as well as any sequences having significant homology to the Pfam and Swissprot protein databases.Finally, the removal of likely erroneous mitochondrial and short transcripts was done by a homemade script.

### Transcriptome Comparisons

Comparisons of our refined transcriptome to the four public horse transcriptomes were done using Cuffcompare [15]. In any pairwise comparison, two transcripts are considered matching if they have the exact intron chain, despite differing terminal exons (class code “=”). If the transcripts are not matching but sharing one or more splice junctions (class code “j”), these would be considered similar transcripts. A transcript is considered novel if it does not overlap with any gene model in the 2^nd^ reference assembly (class code “u”). All other class codes including any kind of overlap with a reference annotation on the opposite strand were considered as “other”. For more detailed descriptions of the class codes provided by Cuffcompare, please see their manual [15]. Complex loci were flagged if a gene model of one assembly overlapped with 2 different gene models in the other assembly. Sensitivity and specificity relative to a given reference transcriptome were calculated per base, intron and locus for each transcriptome and reference combination as described by Burset and Guigó [14].

### Novel Gene Prediction

Any transcript in our final *refined* transcriptome is defined as novel if it does not overlap with a gene model in at least one of the two public equine assemblies, NCBI and ENSEMBL (Cuffcompare class code “u”). Transcripts considered novel were divided into three groups according to the degree of supportive evidence. Transcripts novel to either the NCBI or ENSEMBL assemblies with transcriptional supportive evidence from the other or any other public assembly [5, 7] were in the first category of novel transcripts. Supportive evidence is defined as any overlapping with exon sequence (Cuffcompare class code “=”,“j”,”o”,”x” or “c”). The second and third categories of novel transcripts required that the transcript be absent in all current equine transcriptomes. Transcripts in the second category have supportive evidences in non-horse alignment gene models or *ab initio* gene prediction tracks from the UCSC genome browser. The third category of novel transcripts included transcripts that lack such evidence but have ORFs.

### Tissue-Specific Characterization of The Transcriptome

Tissue-specific transcriptomes were generated by back-mapping the input trimmed RNA-seq reads with Salmon [30] to the refined version of the transcriptome to obtain expression information on a tissue-specific level. A transcript is considered expressed in a given tissue if it has a TPM more than 5% of the total TPM per locus calculated from the tissue specific libraries only. Biological processes identified within the tissue-specific gene blocks were annotated with Panther [32] and reported if the p-values were below the Bonferroni-corrected threshold (5% experiment-wide).

### UCSC Track hubs

Gene Annotation Format (GTF) files were converted into the binary bigbed files [33] using UCSC kintUtils (https://github.com/ENCODE-DCC/kentUtils). The track hub directory structure was designed as recommend by UCSC genome browser [34]. Tracks were constructed using “bigBed 12” format and multiple libraries of the same tissue were organized in composite tracks. The hub files are hosted on a github server as a part of the horse_trans repository (https://github.com/dib-lab/horse_trans).

## List of Abbreviations

(SNP): Single Nucleotide polymorphism
(UTR): Untranslated region
(CNS): Central nervous system
(ICM): Inner cell mass
(TE): Trophectoderm
(RNA-seq): RNA sequencing

## Declarations

### Ethics approval and consent to participate

Not applicable

### Consent for publication

All consent for publication is available

### Availability of data and materials

The data including the scripts for the pipeline as well as the GTF files for the transcriptome can be found at: https://github.com/dib-lab/horse_trans. (This repository is archived by Zenodo at 10.5281/zenodo.56934). All sequencing reads used in this study have been submitted to NCBI Sequence Read Archive; SRA SRP082284 for muscle samples, SRP073514 for brainstem, SRP073514 and SRP082291 for spinal cord, SRP082342 for cerebellum, ERP001525 for retina, SRP031504 and SRP082454 for embryonic tissues and ERP001524, ERP001525 and ERP005568 for skin.

### Authors ’ Contributions

All authors contributed to study conception and design. TAM produced the annotation pipeline and did the data analysis with oversight on writing of the manuscript as well as figure formation. EYS wrote the manuscript and made the figures. CTB, CJF and JDM aided with experimental design and supervised the whole project. RRB and MJM provided retina and skin data. SJV provided muscle data.PJR provided both embryonic tissue data. CJF provided spinal cord and brainstem data. EYS, JDM and MCP provided the cerebellar RNA-seq data. All authors reviewed and approved the final version of the manuscript.

## Acknowledgements

We would like to thank the Arabian horse foundation and Henry Jastro Shields awards for work done on the cerebellum. For the embryo work, support was given by Centre for Equine Health, UC Davis. University of Minnesota Equine Center supported the muscle transcriptomic work used for this study. The RNA seq data from skin and retina was supported in part by generous donations by Appaloosa breeders who belong to the Appaloosa Project’s Electronic Classroom and by the L. David Dube and Heather Ryan Veterinary HealthResearch Fund from the University of Saskatchewan. For J.D.M., M.J.M. and P.J.R. support was provided by UC Davis Agriculture Experiment Station.

## References

1 Brown JB, Boley N, Eisman R, May GE, Stoiber MH, Duff MO, Booth BW, Wen J, Park S,Suzuki AM et al.: Diversity and dynamics of the Drosophila transcriptome. Nature 2014,512(7515):393–399.

2 Mele M, Ferreira PG,Reverter F, DeLuca DS, Monlong J,SammethM, Young TR, GoldmannJM, Pervouchine DD, Sullivan TJ et al.:Human genomics. The human transcriptome across tissues and individuals. Science 2015, 348(6235):660–665.

3 Okazaki Y, Furuno M, Kasukawa T, Adachi J, Bono H, Kondo S, Nikaido I, Osato N, Saito R,Suzuki H et al.: Analysis of the mouse transcriptome based on functional annotation of 60,770 full-length cDNAs. Nature 2002,420(6915):563–573.

4 Holl HM, Gao S, Fei Z, Andrews C, Brooks SA: Generation of a de novo transcriptome from equine lamellar tissue. BMC Genomics 2015,16(1):739.

5 Pacholewska A, Drogemuller M, Klukowska-Rotzler J, Lanz S, Hamza E, Dermitzakis ET,Marti E, Gerber V, Leeb T, Jagannathan V: The transcriptome of equine peripheral blood mononuclear cells. PLoS One 2015,10(3):e0122011.

6 Coleman SJ, Zeng Z, Wang K, Luo S, Khrebtukova I, Mienaltowski MJ, Schroth GP, Liu J,MacLeod JN: Structural annotation of equine protein-coding genes determined bymRNA sequencing. Anim Genet 2010, 41 Suppl 2:121–130.

7 Hestand MS, Kalbfleisch TS, Coleman SJ, Zeng Z, Liu J, Orlando L, MacLeod JN: Annotation of the Protein Coding Regions of the Equine Genome. PLoS One 2015,10(6):e0124375.

8 Yandell M, Ence D: A beginner's guide to eukaryotic genome annotation. Nat Rev Genet2012,13(5):329–342.

9 Curwen V, Eyras E, Andrews TD, Clarke L, Mongin E, Searle SM, Clamp M: The Ensembl automatic gene annotation system. Genome Res 2004,14(5):942–950.

10 Kitts P: The NCBI Handbook: Chapter 14. Genome Assembly and Annotation Process. In: National Center for Biotechnology. 2003 edn: McEntyre, J. & Ostell, J.; 2003.

11 Taanman JW: The mitochondrial genome: structure, transcription, translation and replication. Biochim Biophys Acta 1999,1410(2):103–123.

12 Plotz G, Casper M, Raedle J, Hinrichsen I, Heckel V, Brieger A, Trojan J, Zeuzem S: MUTYH gene expression and alternative splicing in controls and polyposis patients. Hum Mutat 2012, 33(7):1067–1074.

13 Finno CJ, Bordbari M, Monsour T, Bannasch DL, Mickelson DB, Valberg SJ: Spinal cord transcriptome profiling of equine vitamin E deficient neuroaxonal dystrophy identifies dysregulation of cholesterol homeostasis with upregulation of liver X receptor target genes. In Review, Free Rad Biol Med 2016.

14 Burset M, Guigo R: Evaluation of gene structure prediction programs. Genomics 1996,34(3):353–367.

15 Trapnell C, Hendrickson DG, Sauvageau M, Goff L, Rinn JL, Pachter L: Differential analysis of gene regulation at transcript resolution with RNA-seq. Nat Biotechnol 2013,31(l):46–53.

16 Bertone P, Stolc V, Royce TE, Rozowsky JS, Urban AE, Zhu X, Rinn JL, Tongprasit W, Samanta M, Weissman S et al.: Global identification of human transcribed sequences with genome tiling arrays. Science 2004, 306(5705):2242–2246.

17 Khaitovich P, Kelso J, Franz H, Visagie J, Giger T, Joerchel S, Petzold E, Green RE, Lachmann M, Paabo S: Functionality of intergenic transcription: an evolutionary comparison. PLoS Genet 2006, 2(10):el71.

18 Schadt EE, Edwards SW, GuhaThakurta D, Holder D, Ying L, Svetnik V, Leonardson A, Hart KW, Russell A, Li G et al.: A comprehensive transcript index of the human genome generated using microarrays and computational approaches. Genome Biol 2004,5(10):R73.

19 Sultan M, Amstislavskiy V, Risch T, Schuette M, Dokel S, Ralser M, Balzereit D, Lehrach H,Yaspo ML: Influence of RNA extraction methods and library selection schemes on RNA-seq data. BMC Genomics 2014,15:675.

20 Smibert P, Miura P, Westholm JO, Shenker S, May G, Duff MO, Zhang D, Eads BD, Carlson J,Brown JB et al.: Global patterns of tissue-specific alternative polyadenylation in Drosophila. Cell Rep 2012, 1(3):277–289,.

21 Mangone M, Manoharan AP, Thierry-Mieg D, Thierry-Mieg J, Han T, Mackowiak SD, Mis E, Zegar C, Gutwein MR, Khivansara V et al.: The landscape of C. elegans 3'UTRs. Science 2010, 329(5990):432–435.

22 Ulitsky I, Shkumatava A, Jan CH, Subtelny AO, Koppstein D, Bell GW, Sive H, Bartel DP:Extensive alternative polyadenylation during zebrafish development. Genome Res2012, 22(10):2054–2066.

23 Scott EY, Penedo MC, Murray JD, Finno CJ: Defining Trends in Global Gene Expression in Arabian Horses with Cerebellar Abiotrophy. In Review, Cerebellum 2016.

24 Wu JQ, Habegger L, Noisa P, Szekely A, Qiu C, Hutchison S, Raha D, Egholm M, Lin H,Weissman S et al.: Dynamic transcriptomes during neural differentiation of human embryonic stem cells revealed by short, long, and paired-end sequencing. Proc Natl AcadSci U S A 2010, 107(ll):5254–5259.

25 Farkas MH, Grant GR, White JA, Sousa ME, Consugar MB, Pierce EA: Transcriptome analyses of the human retina identify unprecedented transcript diversity and 3.5 Mb of novel transcribed sequence via significant alternative splicing and novel genes. BMC Genomics 2013,14:486.

26 Bolger AM, Lohse M, Usadel B: Trimmomatic: a flexible trimmer for Illumina sequence data. Bioinformatics 2014, 30(15):2114–2120.

27 Macmanes MD: On the optimal trimming of high-throughput mRNA sequence data.Front Genet 2014, 5:13.

28 Kim D, Pertea G, Trapnell C, Pimentel H, Kelley R, Salzberg SL: TopHat2: accurate alignment of transcriptomes in the presence of insertions, deletions and gene fusions. Genome Biol 2013,14(4):R36.

29 Brown C, Howe, A., Zhang, Q., Pyrkosz, A.B. & Brom, T.H.: A Reference-Free Algorithm for Computational Normalization of Shotgun Sequencing Data. 2013.

30 Rob Patro GD, Carl Kingsford: Accurate, fast, and model-aware transcript expression quanitification with Salmon. bioRxiv 2015.

31 Haas BJ, Papanicolaou A, Yassour M, Grabherr M, Blood PD, Bowden J, Couger MB, Eccles D,Li B, Lieber M et al.: De novo transcript sequence reconstruction from RNA-seq using the Trinity platform for reference generation and analysis. Nat Protoc 2013,8(8):1494–1512.

32 Mi H, Muruganujan A, Casagrande JT, Thomas PD: Large-scale gene function analysis with the PANTHER classification system. Nat Protoc 2013, 8(8):1551–1566.

33 Kent WJ, Zweig AS, Barber G, Hinrichs AS, Karolchik D: BigWig and BigBed: enabling browsing of large distributed datasets. Bioinformatics 2010, 26(17):2204–2207.

34 Raney BJ, Dreszer TR, Barber GP, Clawson H, Fujita PA, Wang T, Nguyen N, Paten B, ZweigAS, Karolchik D et al.: Track data hubs enable visualization of user-defined genome-wide annotations on the UCSC Genome Browser. Bioinformatics 2014, 30(7):1003–1005.

